# Traditional Physical Practice Participation and Vision-Related Quality of Life in Adolescents: The Serial Mediating Roles of Exercise Self-Efficacy and Visual Function Anomalies

**DOI:** 10.64898/2026.04.04.716449

**Authors:** Xinxin Zhang, Zhenyu Liu, Jin Long

## Abstract

**Purpose:** This study examined the association between traditional physical practice participation and vision-related quality of life among junior secondary school students and tested the mediating roles of exercise self-efficacy and visual function anomalies within a serial mediation framework.

**Methods:** A four-wave time-lagged survey was conducted among 1,579 students in Grades 7–9 from schools implementing traditional physical practice activities. Variables were assessed at two-week intervals. Mediation effects were tested using the bias-corrected percentile bootstrap method with 5,000 resamples.

**Results:** The total effect of traditional physical practice participation on vision-related quality of life was significant (β = 0.591, p < .001). After including the mediators, the direct effect remained significant (β = 0.404, 95% CI [0.348, 0.457]), accounting for 68.36% of the total effect. The total indirect effect was significant (β = 0.187, 95% CI [0.160, 0.218]), representing 31.64% of the total effect. The indirect effect via exercise self-efficacy was significant (β = 0.088, 95% CI [0.068, 0.112], 14.89%), as was the indirect effect via visual function anomalies (β = 0.065, 95% CI [0.048, 0.086], 11.00%). The serial mediation pathway through exercise self-efficacy and visual function anomalies was also significant (β = 0.034, 95% CI [0.025, 0.045], 5.75%). All confidence intervals excluded zero, supporting partial mediation.

**Conclusion:** Traditional physical practice participation was associated with vision-related quality of life both directly and indirectly through exercise self-efficacy and visual function anomalies, including a significant serial mediation pathway. The findings highlight the combined psychological and functional mechanisms underlying adolescents’ vision-related quality of life.

## 1. Introduction

Visual health has become a salient developmental and public-health concern during adolescence, a period marked by intensive near-work demands, rapidly increasing digital exposure, and high academic load. Global syntheses indicate that myopia already affects a large proportion of children and adolescents, with pooled estimates around one-third and projections pointing to several hundred million affected youths by mid-century (Liang et al., 2025). Beyond refractive status, a broader set of functional visual problems frequently emerges in school-age populations, including visual fatigue, headaches, blurred vision, and reduced reading comfort (Junghans et al., 2020). Such symptoms can interfere with sustained learning, limit participation in daily activities, and compromise overall well-being (Junghans et al., 2020).

Vision-related quality of life (VRQOL) captures the lived consequences of visual impairment by emphasizing functional performance and perceived difficulty in everyday contexts rather than clinical indicators alone (Chen et al., 2024; Wu et al., 2024). Instruments developed for vision-specific quality of life measurement, such as the Low Vision Quality-of-Life Questionnaire (LVQOL), explicitly assess domains spanning distance vision and mobility under varied lighting, adaptation to vision-related limitations, reading and fine work, and activities of daily living (Wolffsohn & Cochrane, 2000). Adolescence is a particularly relevant stage for VRQOL research because visual demands are tightly coupled with academic tasks that rely on sustained close work and continuous attentional engagement (Wu et al., 2024). Functional symptoms can therefore translate into disproportionate educational and psychosocial costs even when standard acuity measures appear acceptable (Chen et al., 2024).

Among functional visual problems, symptom clusters associated with binocular vision dysfunction and related visual function anomalies have attracted growing attention in educational and behavioral settings (Junghans et al., 2020). The Convergence Insufficiency Symptom Survey (CISS) has been validated as a symptom measure for school-age populations, with evidence supporting a threshold score of 16 or higher to distinguish symptomatic cases in children and adolescents (Rouse et al., 2009; Junghans et al., 2020). Symptoms indexed by CISS, including reading fatigue, headaches, difficulty concentrating, blurred vision, and diplopia, align closely with the task ecology of modern schooling (Junghans et al., 2020). Such symptomatology is plausible as a proximal determinant of VRQOL because discomfort during reading and near work can degrade performance efficiency, increase avoidance, and reshape perceptions of daily functioning (Junghans et al., 2020).

Lifestyle behaviors represent a modifiable leverage point for adolescent visual health. Outdoor exposure has demonstrated protective associations with myopia onset and incidence in controlled and quasi-controlled intervention evidence, supporting the role of behavioral environment in visual development (Cao et al., 2020; Deng & Pang, 2019). A complementary line of evidence has begun to position physical activity and reduced sedentary behavior as potentially relevant to myopia risk and eye-related outcomes; a recent dose–response meta-analytic synthesis reported that increasing physical activity while limiting screen time and near work is associated with lower myopia risk among children and adolescents (Ding et al., 2025; Ha et al., 2025). Even when refractive outcomes remain the dominant endpoint, physical activity may matter for vision-related functioning through pathways linked to eye strain, fatigue, posture, autonomic regulation, and general physiological resilience (Junghans et al., 2020).

Traditional physical practices (often operationalized in East Asian contexts as traditional mind–body exercises and health-oriented martial routines) constitute an especially practical activity option for adolescents because participation can occur in school settings, requires minimal equipment, and often emphasizes controlled breathing, postural alignment, and sustained attention. Evidence in youth populations has primarily focused on psychological and cognitive outcomes, with syntheses suggesting potential benefits of Tai Chi or Qigong-style training for stress-related outcomes in adolescents. Nevertheless, empirical work linking traditional practice participation to adolescent VRQOL remains limited, and mechanisms specific to visual functioning have rarely been examined.

Mechanistic clarification is essential because behavioral exposure alone rarely explains variance in VRQOL without accounting for psychological and functional mediators. Exercise self-efficacy, a domain-specific belief in the capability to initiate and maintain exercise behavior under barriers, is positioned in social-cognitive frameworks as a central determinant of sustained physical activity patterns and related health outcomes (Lin et al., 2024). Youth-focused evidence indicates that physical-activity interventions can meaningfully influence self-efficacy, and adolescent observational findings frequently show positive covariation between physical activity and self-efficacy (Lu et al., 2024). Traditional practices may be particularly potent for self-efficacy formation because training typically provides repeated mastery experiences, structured progression, and salient feedback on bodily control. In parallel, reductions in visual function anomalies constitute a plausible functional pathway linking activity participation to VRQOL, given the direct relevance of visual discomfort to reading performance and daily functioning (Junghans et al., 2020).

A serial mediation perspective offers an integrative framework for understanding how traditional practice participation may translate into improved VRQOL: participation may strengthen exercise self-efficacy, elevated self-efficacy may support more consistent engagement and healthier daily routines, and improved routines may coincide with fewer or less severe visual function anomalies, culminating in better vision-related quality of life (Lin et al., 2024). Such a chain model has practical value for intervention design because it identifies targetable psychological resources and symptom-level outcomes rather than relying solely on distal refractive indicators.

The present research evaluates the association between traditional physical practice participation and adolescent VRQOL while examining exercise self-efficacy and visual function anomalies as sequential mediators. Physical activity level is assessed via PARS-3, exercise self-efficacy via ESES, visual function anomalies via CISS, and VRQOL via LVQOL with its multidimensional structure (Rouse et al., 2009; Wolffsohn & Cochrane, 2000). By positioning symptom burden and self-efficacy within a single explanatory pathway, the analysis aims to enrich understanding of how culturally grounded activity modalities may support adolescent visual functioning and daily-life adaptation, offering evidence relevant to school-based health promotion and vision-related well-being.

## 2. Literature review and hypotheses

### 2.1 Direct association between traditional physical practice participation and vision-related quality of life

Traditional physical practices (e.g., Tai Chi- or Qigong-like routines, health-oriented martial forms) are commonly described as mind–body exercise modalities characterized by coordinated movement sequences, controlled breathing, postural alignment, and sustained attentional engagement. In adolescent contexts, such practices are often implemented as low-cost, low-barrier activities that can be integrated into school-based or community-based routines, making participation feasible even under academic time pressure.

Vision-related quality of life (VRQOL) concerns perceived functioning and difficulty in daily life under visual demands, rather than ophthalmic indicators alone (Chen et al., 2024; Wu et al., 2024). For adolescents, the daily-life ecology is heavily shaped by near-work tasks (reading, writing, screen-based learning) and prolonged attentional requirements (Ding et al., 2025; Ha et al., 2025). Under these conditions, VRQOL is likely to reflect not only visual clarity but also comfort, task endurance, mobility under lighting variation, and perceived competence in everyday activities (Wolffsohn & Cochrane, 2000). Participation in traditional physical practices is plausibly relevant to VRQOL because such participation is frequently accompanied by broader patterns of health-related behavior, including structured daily routines, greater engagement in physical activity, and better perceived physical condition (Lin et al., 2024). Prior behavioral health research has repeatedly suggested that sustained participation in organized or structured activity tends to co-vary with quality-of-life indicators in youth, providing an empirical basis for expecting a positive association at the level of VRQOL (Lin et al., 2024).

### 2.2 Indirect association via exercise self-efficacy

Exercise self-efficacy refers to confidence in maintaining physical activity despite barriers such as fatigue, negative mood, lack of support, or limited facilities (Lin et al., 2024). Social-cognitive perspectives treat efficacy beliefs as central correlates of behavioral persistence and relapse recovery (Lin et al., 2024). Traditional physical practices typically involve repeated rehearsal of standardized routines and observable incremental progress in balance, coordination, and movement control. Such training features align closely with mastery experiences and self-regulatory feedback, two conditions frequently discussed as supportive of stronger efficacy beliefs (Lin et al., 2024).

Exercise self-efficacy is also relevant to quality-of-life outcomes because efficacy beliefs often covary with perceived control over health-related routines and coping under stress (Lin et al., 2024). In adolescence, stronger exercise self-efficacy has been linked in many studies to more stable physical activity patterns and better self-management in demanding environments (Lin et al., 2024). VRQOL may be higher when daily routines include regular breaks, balanced activity-rest cycles, and lower perceived strain during visually intensive tasks (Junghans et al., 2020). From an associational standpoint, exercise self-efficacy can be positioned as an intermediate psychological factor connecting participation in traditional physical practices to VRQOL differences (Lin et al., 2024).

### 2.3 Indirect association via visual function anomalies

Visual function anomalies, operationalized as symptom burden during near work, include eye strain, headaches, blurred vision, double vision, sleepiness during reading, and concentration difficulty (Rouse et al., 2009; Junghans et al., 2020). Such symptoms are highly salient for adolescents because they map directly onto core school activities and sustained near-demand tasks (Junghans et al., 2020). When symptom burden is higher, reading efficiency and comfort may be lower, and perceived functioning in daily tasks may be correspondingly less favorable. In vision-specific quality-of-life measures, symptom-related limitations are most likely to register in domains involving reading and fine work, adaptation, and everyday activity performance (Wolffsohn & Cochrane, 2000).

Lifestyle and behavioral factors have often been discussed as correlates of visual symptoms, including patterns of prolonged near work, rest behavior, physical fatigue, and general stress (Ding et al., 2025; Ha et al., 2025). Traditional physical practice participation may co-occur with more health-promoting routines and better perceived bodily condition, which may align with lower reported symptom severity during near work (Junghans et al., 2020). In addition, such practices emphasize posture and controlled movement, which is frequently discussed in relation to musculoskeletal strain and fatigue during reading-related tasks; symptom reporting may therefore differ across participation levels even without assuming a deterministic pathway (Junghans et al., 2020).

### 2.4 Serial indirect association via exercise self-efficacy and visual function anomalies

A serial mediation perspective is useful when psychological resources and functional symptoms are expected to align in a sequential pattern (Lin et al., 2024). Exercise self-efficacy represents a relatively stable, domain-specific belief that can covary with persistence in activity routines and adaptive responses when routines are disrupted (Lin et al., 2024). Visual function anomalies represent a proximal functional burden that is directly experienced during reading and near work. A plausible associative ordering places exercise self-efficacy upstream of day-to-day behavioral stability, with symptom burden downstream of those routines and daily task conditions (Junghans et al., 2020).

Within this logic, higher traditional physical practice participation is expected to align with higher exercise self-efficacy (Lin et al., 2024); higher exercise self-efficacy is expected to align with lower symptom burden; lower symptom burden is expected to align with higher VRQOL (Wolffsohn & Cochrane, 2000). The serial specification does not exclude the possibility of direct covariation between participation and symptom burden, or between exercise self-efficacy and VRQOL, and both paths remain theoretically meaningful in adolescent settings where behavioral and perceptual factors are intertwined.

### 2.5 Theoretical model

Figure 1 presents the proposed theoretical structure integrating a direct association and multiple indirect pathways (see Figure 1). The model specifies: (a) a direct path from traditional physical practice participation to VRQOL (H1), (b) a simple indirect path via exercise self-efficacy (H2), (c) a simple indirect path via visual function anomalies (H3), and (d) a serial indirect path in which exercise self-efficacy and visual function anomalies are ordered sequentially (H4). This structure is consistent with a view that adolescent VRQOL is patterned by both psychological resources related to exercise behavior and symptom-level experiences during near work (Junghans et al., 2020).

**Figure 1.**
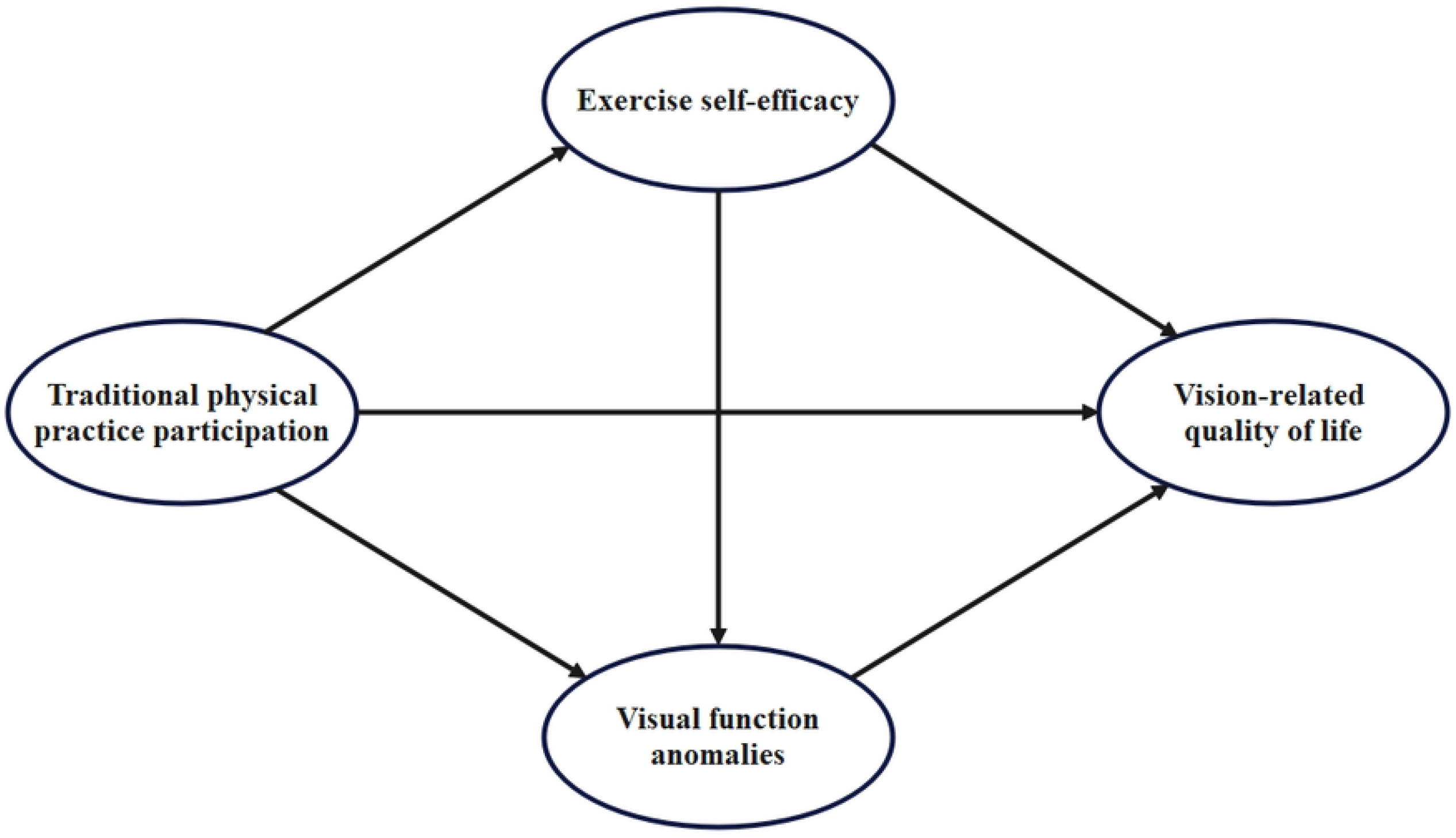
Theoretical model of traditional physical practice participation and vision-related quality of life: A serial mediation framework involving exercise self-efficacy and visual function anomalies.

### 2.6 Hypotheses

**H1:** Traditional physical practice participation is positively associated with vision-related quality of life.

**H2:** Exercise self-efficacy accounts for an indirect association between traditional physical practice participation and vision-related quality of life.

**H3:** Visual function anomalies account for an indirect association between traditional physical practice participation and vision-related quality of life.

**H4:** Traditional physical practice participation shows a serial indirect association with vision-related quality of life via exercise self-efficacy followed by visual function anomalies.

## 3. Materials and methods

### 3.1 Participants and data

A longitudinal survey design with four measurement waves was employed among adolescents in Grades 7–9 (junior secondary school students). Participants were recruited from schools that had implemented traditional physical practice programs as part of regular campus activities, including morning exercise sessions, class-break exercises, after-school activities, or student club programs. These practices primarily involved structured traditional mind–body routines delivered in organized school settings. The first wave was administered between September 1 and September 10, 2025, during which traditional physical practice participation was assessed. The remaining variables were measured at two-week intervals: exercise self-efficacy at Wave 2, visual function anomalies at Wave 3, and vision-related quality of life at Wave 4. A two-week interval was selected to reduce common-method variance and short-term response carryover while preserving relative stability of psychological beliefs and visual symptom experiences during early adolescence. This interval length is frequently adopted in short-term longitudinal behavioral research because it provides temporal separation without substantially increasing attrition. A total of 2,000 questionnaires were distributed using cluster sampling procedures. After excluding cases with patterned responses, substantial missing data, logical inconsistencies, or incomplete participation across waves, 1,579 valid cases were retained, yielding an effective response rate of 78.95%.

Sample size adequacy was evaluated using the item-to-sample ratio heuristic commonly applied in multivariate and structural equation modeling research. An empirical guideline recommends a minimum of 15–20 participants per observed item. The present study included 53 items in total (PARS-3: 3 items; ESES: 10 items; CISS: 15 items; LVQOL: 25 items), resulting in a recommended sample size range of 795 to 1,060 participants. The final valid sample of 1,579 exceeded this recommendation, indicating adequate statistical power and stable parameter estimation for the proposed mediation analyses.

This study was approved by the Ethics Committee of the School of Physical Education, Hainan Normal University (Approval No. HNSFTYXY2026032501). All methods were carried out in accordance with relevant guidelines and regulations, including the principles of the Declaration of Helsinki. Because all participants were minors, written informed consent for study participation was obtained from their parents and/or legal guardians prior to data collection. In addition, written informed consent was also obtained from the adolescent participants themselves before questionnaire administration. Participation was voluntary, and participants had the right to withdraw from the study at any time without penalty.

### 3.2 Measurement tools

All variables were assessed using standardized self-report questionnaires. To maintain comparability across waves and facilitate structural modeling, all instruments were administered using a five-point Likert response format in the present study.

#### Traditional physical practice participation

Participation in traditional physical practices (e.g., school-based Tai Chi, health-oriented martial routines, and structured mind– body exercise programs implemented during class breaks, after-school activities, or student clubs) was assessed using the Physical Activity Rating Scale (PARS-3; Liang, 1994). The scale consists of three items measuring exercise intensity, duration, and frequency. In this study, the mean of the three items was computed to represent the overall level of traditional physical practice participation, with higher mean scores indicating greater participation. A sample item is: “What is the intensity of your usual physical exercise?” Responses range from 1 (very light intensity) to 5 (very vigorous intensity).

#### Exercise self-efficacy

Exercise self-efficacy was measured using the Exercise Self-Efficacy Scale (ESES; Kroll et al., 2007), a 10-item instrument assessing confidence in maintaining physical activity under various barriers. The scale captures general exercise-related efficacy beliefs rather than situation-specific responses to a particular program. Higher mean scores indicate stronger perceived capability to sustain physical activity. A representative item is: “I am confident that I can be physically active or exercise even when I am tired.” Responses range from 1 (strongly disagree) to 5 (strongly agree).

#### Visual function anomalies

Visual function anomalies were assessed using the Convergence Insufficiency Symptom Survey (CISS; Rouse et al., 1999; Convergence Insufficiency Treatment Trial [CITT] Investigator Group, 2009), a 15-item single-factor scale measuring the frequency of visual discomfort during reading or near work. The instrument evaluates current symptom experiences rather than program-specific effects. Higher mean scores indicate greater visual symptom burden. A sample item is: “Do your eyes feel tired when reading or doing close work?” Responses range from 1 (never) to 5 (always).

#### Vision-related quality of life

Vision-related quality of life was measured using the Low Vision Quality of Life Questionnaire (LVQOL; Wolffsohn & Cochrane, 2000), which contains 25 items across four dimensions: Distance Vision, Mobility and Lighting; Adjustment; Reading and Fine Work; and Activities of Daily Living. The scale assesses perceived functional status in vision-related daily tasks. Mean scores were calculated for each dimension and for the overall scale, with higher scores reflecting better perceived functioning. A representative item from the Reading and Fine Work dimension is: “How much difficulty do you have reading small print?” Responses range from 1 (extreme difficulty) to 5 (no difficulty).

### 3.3 Data analysis

All statistical analyses were conducted using IBM SPSS Statistics 26.0 and AMOS 26.0. Prior to hypothesis testing, data screening procedures were performed to examine missing values, outliers, and normality assumptions. Cases with excessive missing data or abnormal response patterns had already been removed during data cleaning. Skewness and kurtosis values were within acceptable ranges, supporting the use of parametric analyses.

Descriptive statistics were computed for all study variables. Internal consistency reliability was assessed using Cronbach’s α, with values above 0.70 indicating acceptable reliability. Composite reliability (CR) and average variance extracted (AVE) were calculated to evaluate construct reliability and convergent validity. Convergent validity was considered adequate when CR > 0.70, AVE > 0.50, and standardized factor loadings exceeded 0.60.

Confirmatory factor analysis (CFA) was conducted to examine the measurement model comprising four latent variables: traditional physical practice participation, exercise self-efficacy, visual function anomalies, and vision-related quality of life. Model fit was evaluated using multiple indices, including the chi-square to degrees of freedom ratio (χ^2^/df), Comparative Fit Index (CFI), Tucker–Lewis Index (TLI), Standardized Root Mean Square Residual (SRMR), and Root Mean Square Error of Approximation (RMSEA) with its 90% confidence interval. Competing measurement models (one-factor, two-factor, three-factor, and four-factor models) were compared to assess discriminant validity.

Given that all constructs were assessed using self-report measures, common method bias (CMB) was examined using both exploratory and confirmatory approaches. Harman’s single-factor test was first conducted using unrotated exploratory factor analysis to determine whether a single factor accounted for the majority of the covariance. Subsequently, CFA-based comparisons of alternative measurement models were performed. In addition, an unmeasured latent method construct (ULMC) model was estimated by including a latent common method factor loading on all observed indicators. CMB was considered negligible when the first unrotated factor explained less than 40% of the total variance and when the ULMC model did not demonstrate a substantially improved fit relative to the hypothesized four-factor model.

Pearson correlation analyses were conducted to examine bivariate associations among the four constructs. Hierarchical multiple regression analyses were then performed to examine the associations between traditional physical practice participation and vision-related quality of life while controlling for gender, grade, and place of residence. Exercise self-efficacy and visual function anomalies were subsequently entered to assess incremental explanatory power and to provide preliminary evidence consistent with mediation. Variance inflation factor (VIF) values were inspected to assess multicollinearity.

To test the hypothesized serial mediation framework, structural equation modeling (SEM) was conducted in AMOS 26.0, specifying both direct and indirect paths from traditional physical practice participation to vision-related quality of life through exercise self-efficacy and visual function anomalies. The structural model was evaluated using the same fit indices described above.

Indirect effects were examined using the bias-corrected percentile bootstrap method with 5,000 resamples. Mediation effects were considered statistically significant when the 95% confidence interval of the bootstrapped estimates did not include zero.

## 4. Result

### 4.1 Sample characteristics

Table 1 presents the demographic characteristics of the final sample (N = 1,579). The gender distribution was balanced, with 797 males (50.47%) and 782 females (49.53%). Participants were evenly distributed across grades, including 515 students in Grade 7 (32.62%), 537 in Grade 8 (34.01%), and 527 in Grade 9 (33.38%). In terms of place of residence, 786 students (49.78%) were from rural or township areas, while 793 students (50.22%) resided in urban areas. The distribution across gender, grade level, and residential background indicates a well-balanced sample of junior secondary school students.

### 4.2 Descriptive statistics and measurement reliability

Descriptive statistics and reliability indices for all study variables are presented in **Table 2**. Internal consistency reliability was satisfactory across all constructs. Cronbach’s α was 0.827 for traditional physical practice participation, 0.938 for exercise self-efficacy, and 0.958 for visual function anomalies. For vision-related quality of life, Cronbach’s α coefficients across dimensions ranged from 0.799 to 0.939, indicating acceptable to excellent reliability.

**Table 1.** Demographic characteristics of the sample (N = 1,579).

| Variable | Category | Frequency | Percentage (%) |
| --- | --- | --- | --- |
| Gender | Male | 797 | 50.47 |
|  | Female | 782 | 49.53 |
| Grade | Grade 7 | 515 | 32.62 |
|  | Grade 8 | 537 | 34.01 |
|  | Grade 9 | 527 | 33.38 |
| Place of residence | Rural/Township | 786 | 49.78 |
|  | Urban | 793 | 50.22 |

**Table 2.**
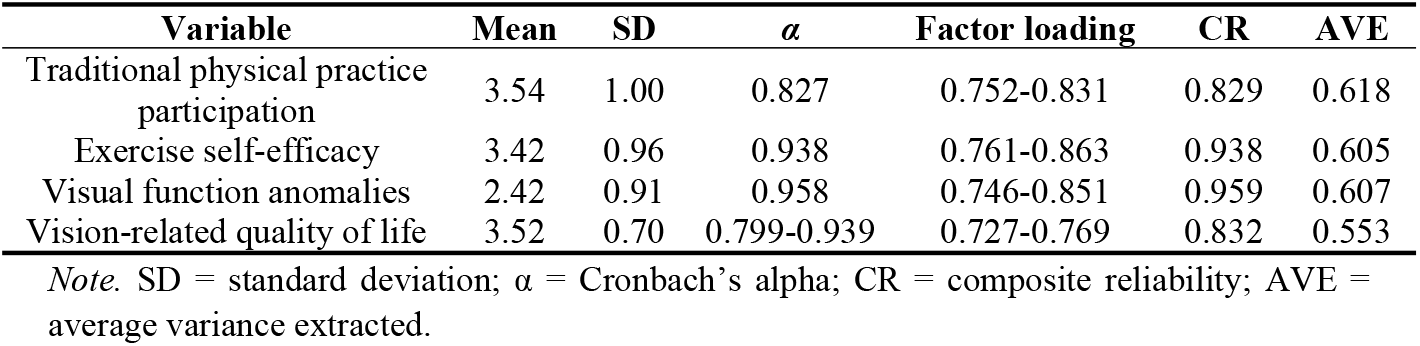
Descriptive Statistics and Measurement Reliability of the Study Variables (N = 1,579).

| Variable | Mean | SD | $\alpha$ | Factor loading | CR | AVE |
| --- | --- | --- | --- | --- | --- | --- |
| Traditional physical practice participation | 3.54 | 1.00 | 0.827 | 0.752-0.831 | 0.829 | 0.618 |
| Exercise self-efficacy | 3.42 | 0.96 | 0.938 | 0.761-0.863 | 0.938 | 0.605 |
| Visual function anomalies | 2.42 | 0.91 | 0.958 | 0.746-0.851 | 0.959 | 0.607 |
| Vision-related quality of life | 3.52 | 0.70 | 0.799-0.939 | 0.727-0.769 | 0.832 | 0.553 |

*Note.* SD = standard deviation; $\alpha$ = Cronbach's alpha; CR = composite reliability; AVE = average variance extracted.

Composite reliability (CR) values ranged from 0.829 to 0.959, all exceeding the recommended threshold of 0.70. Average variance extracted (AVE) values ranged from 0.553 to 0.618, surpassing the recommended criterion of 0.50. Standardized factor loadings were between 0.727 and 0.863 across constructs. These results support adequate internal consistency and convergent validity of the measurement model, providing a sound basis for subsequent structural analyses.

### 4.3 Common method bias test

Given that all variables were measured using self-report questionnaires, potential common method bias (CMB) was examined. Harman’s single-factor test was first conducted using unrotated exploratory factor analysis. The results showed that seven factors had eigenvalues greater than 1, jointly explaining 64.00% of the total variance. The first unrotated factor accounted for 32.72% of the variance, which is below the commonly referenced threshold of 40%, indicating that a single factor did not dominate the covariance structure.

To further assess common method bias, confirmatory factor analyses were conducted to compare alternative measurement models, including one-factor, two-factor, three-factor, and the hypothesized four-factor model. In addition, an unmeasured latent method construct (ULMC) model was estimated by adding a latent common method factor loading on all observed indicators. As presented in **Table 3**, the hypothesized four-factor model demonstrated substantially better fit (χ^2^/df = 2.380, CFI = 0.981, TLI = 0.979, SRMR = 0.017, RMSEA = 0.030) compared with the more constrained models. The ULMC model did not show meaningful improvement over the four-factor model, with identical fit indices. These findings suggest that common method bias was unlikely to threaten the validity of the results.

**Table 3.** Comparison of alternative measurement models for common method bias assessment (N = 1,579).

| Model | $\chi^2/\text{df}$ | CFI | TLI | SRMR | RMSEA (90% CI) |
| --- | --- | --- | --- | --- | --- |
| One-factor | 25.273 | 0.659 | 0.636 | 0.132 | 0.124 (0.122, 0.126) |
| Two-factor | 23.209 | 0.689 | 0.667 | 0.130 | 0.119 (0.117, 0.121) |
| Three-factor | 6.358 | 0.925 | 0.920 | 0.065 | 0.058 (0.056, 0.060) |
| Four-factor | 2.380 | 0.981 | 0.979 | 0.017 | 0.030 (0.027, 0.032) |
| ULMC-factor | 2.386 | 0.981 | 0.979 | 0.017 | 0.030 (0.027, 0.032) |

*Note.* ULMC = unmeasured latent method construct; CFI = comparative fit index; TLI = Tucker–Lewis index; SRMR = standardized root mean square residual; RMSEA = root mean square error of approximation; CI = confidence interval.

### 4.4 Correlation analysis and discriminant validity

Pearson correlation coefficients among the four study variables are presented in **Table** Traditional physical practice participation was positively correlated with exercise self-efficacy (r = 0.299, *p* < .001) and vision-related quality of life (r = 0.477, *p* < .001), and negatively correlated with visual function anomalies (r = −0.347, *p* < .001). Exercise self-efficacy was negatively correlated with visual function anomalies (r = −0.453, *p* < .001) and positively correlated with vision-related quality of life (r = 0.442, *p* < .001). Visual function anomalies showed a negative correlation with vision-related quality of life (r = −0.466, *p* < .001). All correlations were statistically significant at the 0.001 level and consistent with the hypothesized directional relationships.

Discriminant validity was evaluated using the Fornell–Larcker criterion. As shown in **Table 4**, the square roots of the average variance extracted (AVE) for each construct (values on the diagonal) were greater than the corresponding inter-construct correlations. This pattern indicates adequate discriminant validity among traditional physical practice participation, exercise self-efficacy, visual function anomalies, and vision-related quality of life.

**Table 4.** Correlations and discriminant validity among key variables (N = 1,579).

| Variable | Traditional physical practice participation | Exercise self-efficacy | Visual function anomalies | Vision-related quality of life |
| --- | --- | --- | --- | --- |
| Traditional physical practice participation | 0.786 |  |  |  |
| Exercise self-efficacy | 0.299*** | 0.778 |  |  |
| Visual function anomalies | -0.347*** | -0.453*** | 0.779 |  |
| Vision-related quality of life | 0.477*** | 0.442*** | -0.466*** | 0.744 |

*Note.* \*\*\* $p < .001$ . Values on the diagonal represent the square roots of the average variance extracted (AVE).

### 4.5 Multiple regression analysis

Hierarchical multiple regression analyses were conducted to examine the associations among traditional physical practice participation, exercise self-efficacy, visual function anomalies, and vision-related quality of life (see Table 5). Gender, grade, and place of residence were entered as control variables in Model 1. The addition of traditional physical practice participation in Model 2 substantially increased the explained variance. In Model 3, exercise self-efficacy and visual function anomalies were included simultaneously.

**Table 5.** Hierarchical multiple regression analysis predicting vision-related quality of life (N = 1,579).

| Variables | Model 1 |  | Model 2 |  | Model 3 |  |
| --- | --- | --- | --- | --- | --- | --- |
| | $\beta$ | SE | $\beta$ | SE | $\beta$ | SE |
| **Control variables** |  |  |  |  |  |  |
| Gender | 0.115**<br>* | 0.035 | 0.073 | 0.031*<br>* | 0.068*** | 0.028 |
| Grade | -0.010 | 0.022 | -0.003 | 0.019 | -0.003 | 0.017 |
| Place of residence | -0.029 | 0.035 | -0.025 | 0.031 | -0.012 | -0.028 |
| **Independent variable** |  |  |  |  |  |  |
| Traditional physical practice participation |  |  | 0.470**<br>* | 0.016 | 0.315*** | 0.015 |
| **Mediator** |  |  |  |  |  |  |
| Exercise self-efficacy |  |  |  |  | 0.232*** | 0.017 |
| Visual function anomalies |  |  |  |  | -0.249**<br>* | 0.018 |
| **Model summary** |  |  |  |  |  |  |
| R<sup>2</sup> |  | 0.014 |  | 0.233 |  | 0.377 |
| $\Delta R^2$ | | 0.014 | | 0.219 | | 0.144 |
| F |  | 7.489*** |  | 119.617*** |  | 158.380*** |

*Note.* Standardized coefficients ( $\beta$ ) are reported. \* $p < .05$ , \*\* $p < .01$ , \*\*\* $p < .001$ .

Traditional physical practice participation showed a significant positive association with vision-related quality of life. After including the mediators, the standardized coefficient for traditional physical practice participation decreased but remained significant, while exercise self-efficacy was positively associated and visual function anomalies were negatively associated with vision-related quality of life. The final model explained 37.7% of the variance. Variance inflation factor (VIF) values were all below 1.5, indicating no multicollinearity concerns. Overall, the pattern of coefficients was consistent with the proposed mediation framework.

### 4.6 Structural equation modeling analysis

Structural equation modeling (SEM) was conducted using AMOS 26.0 to test the hypothesized serial mediation model. The structural model demonstrated good overall fit to the data, with χ^2^/df = 2.380, CFI = 0.981, TLI = 0.979, SRMR = 0.017, and RMSEA = 0.030 (90% CI [0.027, 0.032]). All indices met the recommended criteria (χ^2^/df < 5; CFI and TLI > 0.90; SRMR < 0.08; RMSEA < 0.08), indicating satisfactory model fit.

The standardized path coefficients are illustrated in **Figure 2**, reflecting both direct and indirect associations consistent with the proposed structural framework. All structural paths were statistically significant. Traditional physical practice participation showed a positive direct association with vision-related quality of life (β = 0.40). It was positively associated with exercise self-efficacy (β = 0.34) and negatively associated with visual function anomalies (β = −0.26). Exercise self-efficacy was negatively associated with visual function anomalies (β = −0.39) and positively associated with vision-related quality of life (β = 0.26). Visual function anomalies showed a negative association with vision-related quality of life (β = −0.25).

**Figure 2.**
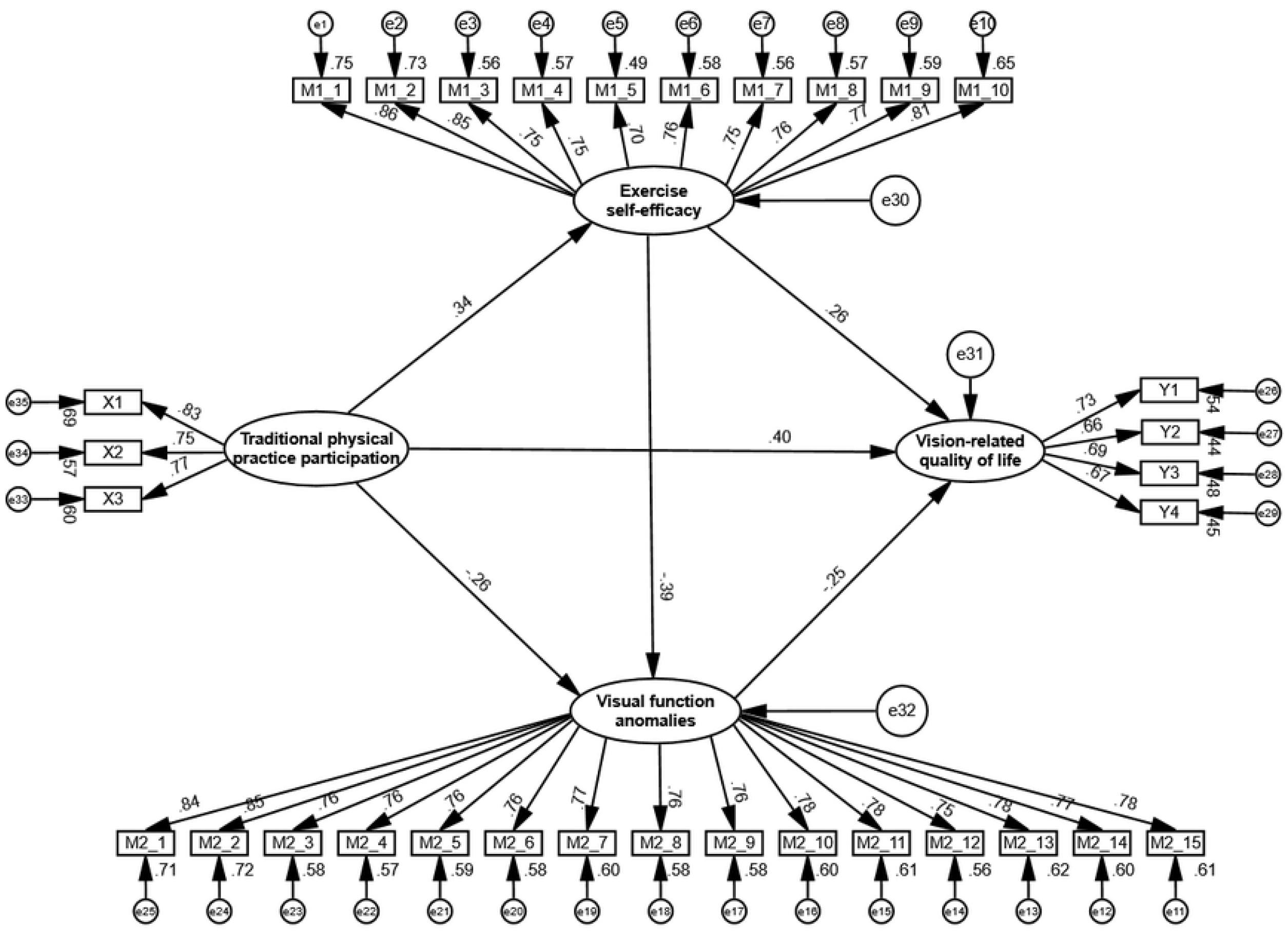
Standardized path coefficients of the structural model.

### 4.7 Bootstrapping mediation test

The mediation effects were examined using the bias-corrected percentile bootstrap method with 5,000 resamples. A mediation effect was considered statistically significant when the 95% confidence interval did not include zero. The results are presented in **Table 6**.

**Table 6.** Bootstrapping Results for Direct and Indirect Effects (N = 1,579).

| Path | $\beta$ | Boot SE | Boot LLCI | Boot ULCI | Ratio (%) |
| --- | --- | --- | --- | --- | --- |
| **Direct effect** | 0.404*** | 0.028 | 0.348 | 0.457 | 68.36 |
| **Indirect effects** | 0.187*** | 0.014 | 0.160 | 0.218 | 31.64 |
| Traditional physical practice participation→Exercise self-efficacy→vision-related quality of life | 0.088*** | 0.011 | 0.068 | 0.112 | 14.89 |
| Traditional physical practice participation→Visual function anomalies→vision-related quality of life | 0.065*** | 0.010 | 0.048 | 0.086 | 11.00 |
| Traditional physical practice participation→Exercise self-efficacy→Visual function anomalies→vision-related quality of life | 0.034*** | 0.005 | 0.025 | 0.045 | 5.75 |
| **Total effect** | 0.591*** | 0.024 | 0.542 | 0.635 | 100 |

Note. \*\*\* $p < .001$ . Boot LLCI, the lower bound of the 95% confidence interval. Boot ULCI, the upper limit of the 95% confidence interval (bias-corrected percentile bootstrap method). Bootstrap sample size = 5,000.

The total effect of traditional physical practice participation on vision-related quality of life was significant (β = 0.591, p < .001). The direct effect remained significant after including the mediators (β = 0.404, 95% CI [0.348, 0.457]), accounting for 68.36% of the total effect. The overall indirect effect was also significant (β = 0.187, 95% CI [0.160, 0.218]), representing 31.64% of the total effect, indicating the presence of partial mediation.

Regarding specific indirect pathways, the indirect effect via exercise self-efficacy was significant (β = 0.088, 95% CI [0.068, 0.112]), accounting for 14.89% of the total effect. The indirect effect via visual function anomalies was also significant (β = 0.065, 95% CI [0.048, 0.086]), accounting for 11.00% of the total effect. The serial mediation pathway through exercise self-efficacy and visual function anomalies was significant as well (β = 0.034, 95% CI [0.025, 0.045]), accounting for 5.75% of the total effect. None of the confidence intervals included zero, supporting the robustness of the mediation effects.

In terms of hypothesis testing, the direct association between traditional physical practice participation and vision-related quality of life was supported (H1). The indirect effect through exercise self-efficacy was supported (H2), and the indirect effect through visual function anomalies was supported (H3). The serial mediation pathway involving exercise self-efficacy followed by visual function anomalies was also supported (H4).

## 5. Discussion

### 5.1 Overview of main findings

The analysis examined associations among traditional physical practice participation, exercise self-efficacy, visual function anomalies, and vision-related quality of life in Grades 7– 9 adolescents, using a four-wave time-lagged design implemented in schools offering traditional practice activities during class breaks, after-school sessions, or club programs. Results supported a positive association between traditional physical practice participation and vision-related quality of life. Exercise self-efficacy and visual function anomalies each accounted for significant indirect associations, and the serial pathway linking exercise self-efficacy to visual function anomalies also accounted for a significant portion of the overall association. The coexistence of a significant direct effect with significant indirect effects was consistent with a partial mediation pattern, suggesting that multiple pathways may co-occur within the proposed framework (Podsakoff et al., 2024; Kock et al., 2021).

### 5.2 Direct association between traditional physical practice participation and vision-related quality of life

Traditional physical practice participation showed a positive direct association with vision-related quality of life after inclusion of the mediators. In early adolescence, vision-related quality of life is strongly shaped by sustained near-work demands, frequent reading-related tasks, and prolonged classroom attention. Traditional physical practice routines delivered in structured school settings often involve regulated breathing, coordinated movement, and postural control, alongside a predictable participation schedule. Such characteristics align with broader patterns of health-related routine formation and perceived functional readiness in daily tasks. A positive association with vision-related quality of life can be interpreted as consistent with a behavioral health perspective in which engagement in structured physical practice co-varies with more favorable perceptions of daily functioning, including mobility under different lighting conditions, adaptation to vision-related constraints, reading-related comfort, and activities of daily living (Dutheil et al., 2023; Ha et al., 2025; Lu et al., 2024; Mataftsi et al., 2023).

The magnitude of the direct association also indicates that exercise self-efficacy and visual function anomalies, while important, do not fully account for the linkage between participation and vision-related quality of life. Additional correlates may contribute, including sleep quality, stress levels, screen-time patterns, near-work scheduling, habitual rest breaks, or general physical fitness. Such factors were not modeled as mediators in the present framework yet remain plausible contributors to the residual direct association (Ha et al., 2025; Dutheil et al., 2023; Jin et al., 2024; Liu, 2023; Datta et al., 2023; Mataftsi et al., 2023).

### 5.3 Exercise self-efficacy pathway

Exercise self-efficacy accounted for a significant indirect association between traditional physical practice participation and vision-related quality of life. Traditional physical practices typically provide repeated mastery experiences through standardized sequences, progressive skill refinement, and visible improvement in balance, coordination, and bodily control. In social-cognitive terms, mastery experiences and perceived progress are closely aligned with efficacy beliefs, particularly in contexts where participation barriers include fatigue, time pressure, and variable peer support. School-based delivery during class breaks, after-school activities, or clubs also reduces structural barriers, allowing efficacy beliefs to develop within a stable routine (Lin et al., 2024; Shan et al., 2026; Moeller et al., 2024).

The presence of this indirect association aligns with a broader view that self-efficacy beliefs help individuals sustain engagement, manage perceived obstacles, and generalize competence beliefs into health-related contexts. Adolescents with higher exercise self-efficacy may perceive greater personal agency in managing physical and attentional demands, which may translate into more favorable evaluations of functional daily performance, including vision-related functioning. The indirect pathway suggests that traditional practice participation may contribute to perceived quality of life partly by strengthening confidence in physical capability and behavioral control (Shan et al., 2026; Lin et al., 2024).

### 5.4 Visual function anomalies pathway

Visual function anomalies accounted for a second significant indirect association between traditional physical practice participation and vision-related quality of life. Visual function anomalies capture symptom experiences during reading or near work, including eye strain, headaches, blurred vision, and concentration difficulty. Such symptoms represent a proximal functional burden with direct relevance to school learning, reading endurance, and daily task efficiency. A negative association between symptom burden and vision-related quality of life aligns with the structure of LVQOL domains, particularly reading and fine work, adjustment, and everyday activity functioning (Kaur et al., 2022; Mataftsi et al., 2023; Şambel Aykutlu et al., 2024; Zou et al., 2026).

The indirect association implies that traditional physical practice participation may co-vary with reduced symptom burden, which in turn aligns with higher perceived vision-related quality of life. Several plausible mechanisms may underlie this pattern. Traditional practices often encourage postural alignment, controlled breathing, and brief periods of physical disengagement from sustained near work. Participation may also increase general activity levels and reduce uninterrupted screen exposure time. Although the present study did not directly measure screen or near-work behaviors, these behavioral patterns are consistent with factors known to be associated with digital eye strain and asthenopic complaints (Datta et al., 2023; Mataftsi et al., 2023; Kaur et al., 2022).

### 5.5 Serial mediation mechanism

The serial indirect pathway was also significant, indicating that traditional physical practice participation was associated with higher exercise self-efficacy, which in turn was associated with lower visual function anomalies, and thereby with higher vision-related quality of life. This sequential structure suggests that efficacy beliefs may influence symptom experiences, potentially through behavioral regulation and coping processes. Adolescents who feel more capable and confident in managing physical activity may also feel more capable of managing health-related behaviors, such as maintaining appropriate posture, taking breaks during near work, or moderating screen exposure, which could reduce symptom experiences linked to near work and digital device use (Şambel Aykutlu et al., 2024; Kaur et al., 2022; Zou et al., 2026).

The serial mediation results are consistent with a multi-stage interpretation in which psychological resources shape behavioral patterns that influence functional symptom outcomes. Exercise self-efficacy may represent a broader belief in capability that generalizes to self-regulatory behaviors in school and study contexts. Although the current data do not allow detailed behavioral testing of this mechanism, the serial pattern offers a plausible pathway linking participation in structured physical practice to vision-related functioning via psychological and symptom-related processes (Liu et al., 2021; Shan et al., 2026).

### 5.6 Theoretical and practical implications

The use of a four-wave time-lagged design strengthens the temporal ordering of constructs compared to cross-sectional designs. This approach helps reduce same-time measurement artifacts and provides greater inferential clarity in examining indirect pathways. The incorporation of statistical diagnostics for common method bias further supports interpretation of associations observed in self-report data, although such procedures cannot fully eliminate method-related concerns. The current results therefore provide evidence consistent with the hypothesized pathways but should be interpreted as associational rather than causal (Podsakoff et al., 2024; Kock et al., 2021; Shahiwala et al., 2024).

School settings provide a pragmatic context for traditional physical practice implementation, particularly when activities are embedded in class-break exercises, after-school sessions, or club programs. The positive association with vision-related quality of life suggests potential value for school health promotion agendas that seek low-cost, scalable activity modalities. Program design may benefit from explicit attention to self-efficacy formation, since efficacy beliefs represented a meaningful pathway. Instructional strategies that emphasize progressive mastery, attainable goals, and feedback on improvement may strengthen efficacy beliefs. In parallel, monitoring visual symptoms during reading and near work may help identify students with elevated symptom burden, since symptom levels showed strong relevance to vision-related quality of life. Combined emphasis on psychological resources and symptom experiences aligns with the serial mediation findings, supporting a dual-focus approach in school-based health education (Moeller et al., 2024; Shan et al., 2026; Huang et al., 2026; Kaur et al., 2022).

### 5.7 Limitations and future directions

Several limitations warrant consideration. All measures relied on self-report, which may introduce reporting biases despite the time-lagged design and the common method bias diagnostics. The design supports temporal separation across constructs but does not establish causal effects; unmeasured confounders may contribute to the observed associations. Variables such as screen time, near-work duration, sleep quality, refractive status, binocular vision clinical indicators, and socioeconomic factors may correlate with both traditional practice participation and the outcome measures. The sample was drawn from schools already offering traditional physical practice activities, which may limit generalizability to contexts without such programs or to regions with different school schedules and health promotion practices (Podsakoff et al., 2024; Kock et al., 2021; Ha et al., 2025; Dutheil et al., 2023; Jin et al., 2024; Huang et al., 2026).

Future work could incorporate objective behavioral measures (accelerometry for activity, digital usage logs for screen exposure), clinical assessments of visual function, and longer follow-up windows to examine stability and developmental change. Experimental or quasi-experimental designs, such as matched comparison schools or randomized class-level programming, would strengthen causal inference. Additional mediators, including sleep, stress, attentional functioning, and habitual break behavior during near work, may further refine the explanatory model (Ha et al., 2025; Mataftsi et al., 2023; Huang et al., 2026).

## 6. Conclusion

The findings indicated that higher traditional physical practice participation was associated with more favorable vision-related quality of life among Grades 7–9 adolescents in schools implementing traditional practice activities. Exercise self-efficacy and visual function anomalies accounted for significant indirect associations, and the serial pathway linking exercise self-efficacy to visual function anomalies provided additional explanatory value. The overall pattern supports a multi-pathway framework in which psychological resources and functional symptom experiences jointly align with adolescents’ perceived vision-related daily functioning.

## Ethics approval and consent to participate

This study was approved by the Ethics Committee of the School of Physical Education, Hainan Normal University (Approval No. HNSFTYXY2026032501). All methods were carried out in accordance with relevant guidelines and regulations. As all participants were minors, written informed consent for participation was obtained from their parents and/or legal guardians prior to data collection. Written informed consent was also obtained from all adolescent participants. Participation was voluntary, and participants had the right to withdraw at any time without penalty.

## Data Availability

The data that support the findings of this study are available from the corresponding author upon reasonable request.

